# A Bioinspired Glycopolymer for Capturing Membrane Proteins in Native-Like Lipid-Bilayer Nanodiscs

**DOI:** 10.1101/2021.03.31.437849

**Authors:** Bartholomäus Danielczak, Marie Rasche, Julia Lenz, Eugenio Pérez Patallo, Sophie Weyrauch, Florian Mahler, Michael Tope Agbadaola, Annette Meister, Jonathan Oyebamiji Babalola, Carolyn Vargas, Cenek Kolar, Sandro Keller

## Abstract

Amphiphilic copolymers that directly extract membrane proteins and lipids from cellular membranes to form nanodiscs combine the advantages of harsher membrane mimics with those of a native-like membrane environment. Among the few commercial polymers that are capable of forming nanodiscs, alternating diisobutylene/maleic acid (DIBMA) copolymers have gained considerable popularity as gentle and UV-transparent alternatives to aromatic polymers. However, their moderate hydrophobicities and high electric charge densities render all existing aliphatic copolymers rather inefficient under near-physiological conditions. Here, we introduce Glyco-DIBMA, a bioinspired glycopolymer that possesses increased hydrophobicity and reduced charge density but nevertheless retains excellent solubility in aqueous solutions. Glyco-DIBMA outperforms established aliphatic copolymers in that it solubilizes lipid vesicles of various compositions much more efficiently, thereby furnishing smaller, more narrowly distributed nanodiscs that preserve a bilayer architecture and exhibit rapid lipid exchange. We demonstrate the superior performance of Glyco-DIBMA in preparative and analytical applications by extracting a broad range of integral membrane proteins from cellular membranes and further by purifying a membrane-embedded voltage-gated K^+^ channel, which was fluorescently labeled and analyzed with the aid of microfluidic diffusional sizing (MDS) directly within native-like lipid-bilayer nano-discs.

## Introduction

The gentle isolation of membrane proteins from cellular membranes is a major challenge^1^ that is increasingly met by the use of nanodisc-forming amphiphilic polymers such as aliphatic diisobutylene/maleic acid (DIBMA, Figure 1a)^2–8^ or aromatic styrene/maleic acid (SMA) copolymers.^3,9–13^ Unlike traditional head-and-tail detergents, which displace the native lipid environment of membrane proteins,^14^ amphiphilic copolymers co-extract a nanoscopic lipid patch from the cellular membrane to form polymer-bounded lipid-bilayer nanodiscs.^12^ Thus, these polymers allow encapsulated membrane proteins to remain embedded in a near-native lipid bilayer during isolation, purification, and investigation. The advantages offered by this approach manifest in superior membrane-protein stability^6,10,13,15–22^ and more native-like ligand interactions^15^ and other protein activities^6,16,19^ as compared with detergent micelles.

**Figure 1:**
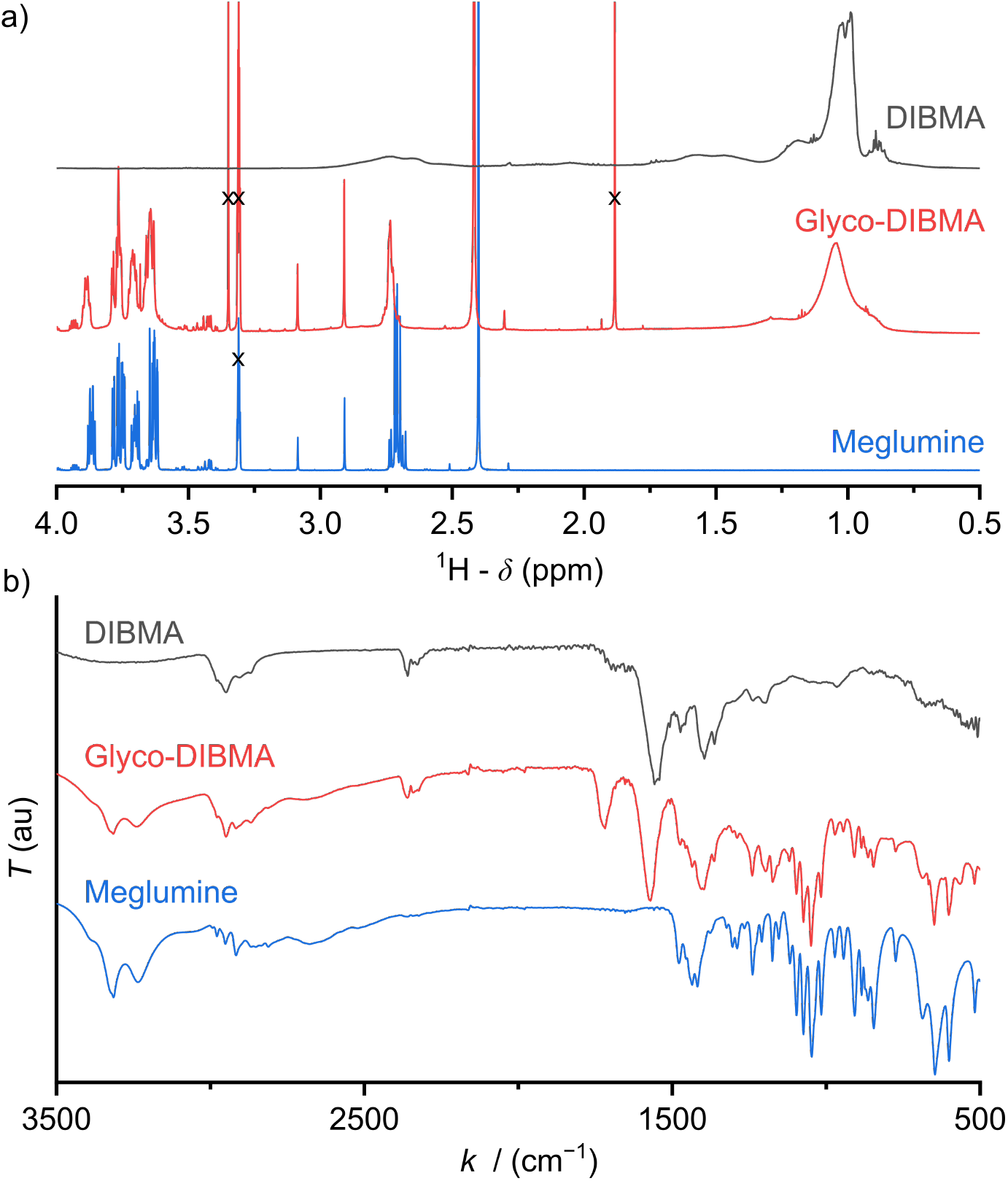
a) ^1^H-NMR spectra of DIBMA, Glyco-DIBMA, and meglumine. The DIBMA spectrum was taken in D_2_O and referenced to the signal of 3-(trimethylsilyl)propane-1-sulfonate (DSS) at 0 ppm, whereas the spectra of Glyco-DIBMA and meglumine were taken in CD_3_OD and referenced to the signal of residual CD_3_OH at 3.31 ppm. Solvent signals are marked with a cross. b) ATR-FTIR spectra of DIBMA, Glyco-DIBMA, and meglumine.

Unlike most other polymers used for extracting membrane proteins,^2,3,8^ DIBMA copolymers contain no aromatic moieties and, thus, have substantially lower background absorption in the UV range. However, these alternating polymers are characterized by a high density of carboxylic acid groups and, consequently, relatively low hydrophobicity. Under physiological conditions of near-neutral pH, the high charge density of DIBMA results in strong Coulombic repulsion between the polymer and anionic lipid membranes on the one hand and among polymer chains on the other.^4,23,24^

These adverse effects reduce the effective membrane affinity and, thereby, the solubilization efficiency of alternating aliphatic polymers. Moreover, Coulombic repulsion is expected to limit the density of polymer chains in the nanodisc rim, which might be responsible for the broad size distributions observed for DIBMA/lipid particles (DIBMALPs).^2,23^ We and others have shown that Coulombic repulsion can be reduced either by adjusting the buffer composition to increase the ionic strength of the solution^2,11,25^ or by reducing the effective charge on the polymer through partial protonation at low pH^23,25^ or through adsorption of divalent cations.^2,4,8,26^ Although effective, all of these remedies render the solution conditions quite unphysiological and, in doing so, partly offset the benefits of alternating aliphatic polymers for handling membrane proteins under conditions that ideally should be as native-like as possible.

On this premise, we sought to develop a new, powerful polymer that is intrinsically less charged and more hydrophobic, with the intention of rendering it an efficient solubilizer also under physiological conditions. At the same time, the new polymer should retain the essential favorable properties of existing aliphatic copolymers and should be synthetically accessible by post-polymerization functionalization of a commercial polymer backbone, thus avoiding the need for laborious and expensive *de novo* polymer synthesis. Herein, we present Glyco-DIBMA (Scheme 1) as a new glycopolymer that is partially amidated with the amino sugar *N*-methyl-D-glucamine (“meglumine”). Utilizing a range of biochemical, biophysical, and imaging techniques, we show that Glyco-DIBMA outperforms established aliphatic copolymers in that it gives rise to smaller and more narrowly distributed lipid-bilayer nanodiscs from both model and cellular membranes under a broad variety of solution conditions. Membrane proteins extracted with the aid of Glyco-DIBMA are amenable to protein purification by chromatography and to further manipulation and analysis under well-controlled yet native-like conditions.

## Results and Discussion

Glycosylation is nature’s most powerful and versatile means of endowing biomacromolecules with enhanced water solubility without accumulating additional electric charge. Inspired by this, we reasoned that glycosylation might provide us with an opportunity to lower the charge density and, thereby, increase the hydrophobicity of a polyanionic polymer backbone without, however, compromising its excellent water solubility. A further crucial requirement was that the synthetic route should be reasonably straightforward and clean such as to afford the final product in sufficient quantity—that is, on the scale of hundreds of grams—and in high purity for compatibility with sensitive biomolecular specimens. In accordance with these considerations, we synthesized Glyco-DIBMA in a two-step process by first converting free DIBMA acid into the more reactive anhydride form and then amidating the latter with meglumine (Scheme 1).

**Scheme 1.**
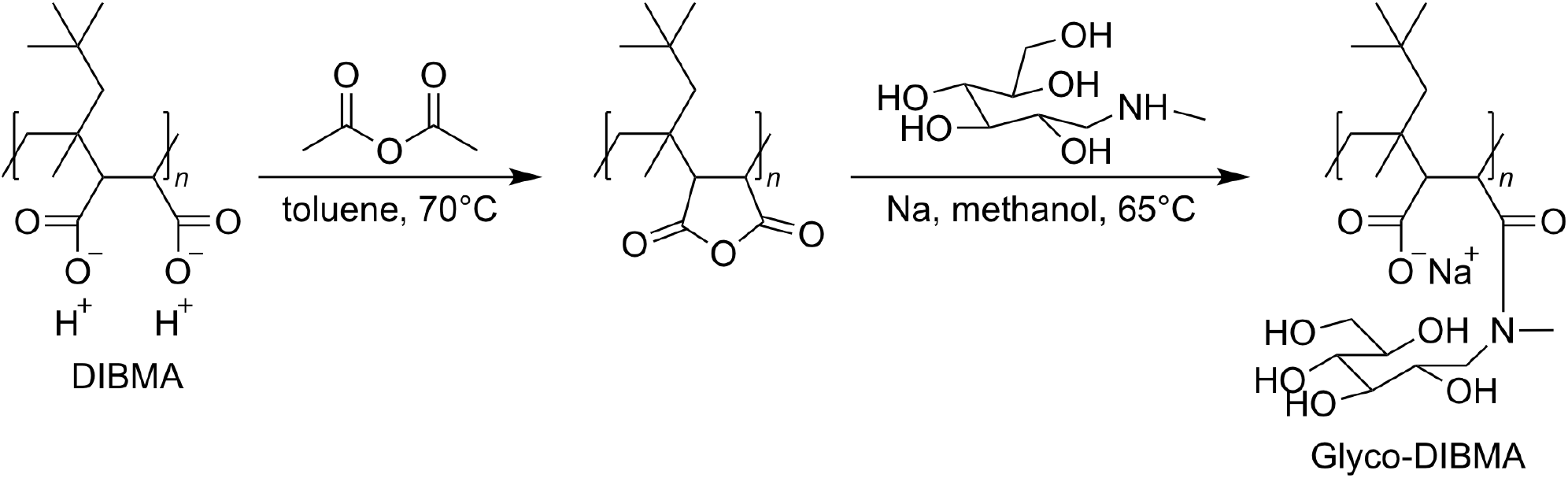
Synthesis of Glyco-DIBMA.

The expected chemical structure of Glyco-DIBMA was corroborated by ^1^H-NMR spectroscopy as well as attenuated total reflectance Fourier-transform infrared (ATR-FTIR) spectroscopy (Figure 1) and its relatively narrow chain length distribution by analytical size exclusion chromatography (SEC) in aqueous phase (Figure SI 1). The ^1^H-NMR spectrum of Glyco-DIBMA displayed the broad polymer peaks also observed for DIBMA and, additionally, the considerably sharper multiplet peaks characteristic of meglumine (Figure 1a). Nevertheless, all signals stemming from the sugar moiety became broader and shifted to lower magnetic field strengths in Glyco-DIBMA as compared with free meglumine (Figure 1a), thus confirming successful coupling. This was further supported by ATR-FTIR (Figure 1b), as the spectrum of Glyco-DIBMA again revealed both the peaks typical of DIBMA and those of meglumine. In SEC (Figure SI 1), Glyco-DIBMA eluted slightly earlier than DIBMA, as expected on the basis of its larger molar mass resulting from glycosylation. However, it should be noted that the elution behavior of polymers also depends on the degree of polymer compaction, that is, the distribution of effective polymer chain extensions in solution. The latter is notoriously difficult to account for quantitatively because it depends not only on intrinsic properties of the polymer such as the number of ionizable groups but also on solution conditions such as the pH value and the concentration of multivalent counterions.^4,23^ In summary, all three methods consistently attested to the high chemical purity and favorable solution behavior of Glyco-DIBMA, thus opening the way for a detailed investigation of its lipid solubilization and protein extraction properties.

We examined the ability of Glyco-DIBMA to spontaneously form lipid-bilayer nanodiscs by mixing it with large unilamellar vesicles (LUVs) made from the zwitterionic phospholipid 1,2-dimyristoyl-*sn*-glycero-3-phosphocholine (DMPC). Dynamic light scattering (DLS) showed that addition of Glyco-DIBMA to DMPC at a mass ratio of polymer (P) to lipid (L) of *m*_P_/*m*_L_ = 1.5 completely abolished the original vesicular structures, which gave way to nanoscopic particles having a *z*-average hydrodynamic diameter and an associated size distribution width of *d_z_ =* (15±6) nm (Figure 2a). These Glyco-DIBMA/lipid particles (Glyco-DIBMALPs) were considerably smaller and more narrowly distributed than DIBMALPs produced at the same mass ratio, which were characterized by *d_z_ =* (31±13) nm. Upon titrating DMPC LUVs with Glyco-DIBMA (Figure 2b), we observed an initial increase in *d_z_* at *m*_P_/*m*_L_ ≤ 0.5 followed by a smooth decrease to reach *d_z_ =* (12±5) nm at *m*_P_/*m*_L_ = 2.5. DIBMA exhibited a qualitatively similar pattern but, throughout the titration, gave rise to larger nanoparticles that displayed broader size distributions and that co-existed with “free” polymer not associated with nanodiscs. Both of these phenomena manifested particularly clearly in analytical SEC, which at *m*_P_/*m*_L_ = 1.5 (Figure 2c) revealed two broad peaks centered at elution volumes of *V*_E_ ≈ 12 mL and 18 mL corresponding to nanodiscs and free polymer, respectively. In stark contrast with this, Glyco-DIBMALPs eluted in a single, sharp peak at *V*_E_ ≈ 17 mL without a significant population of free polymer.

**Figure 2:**
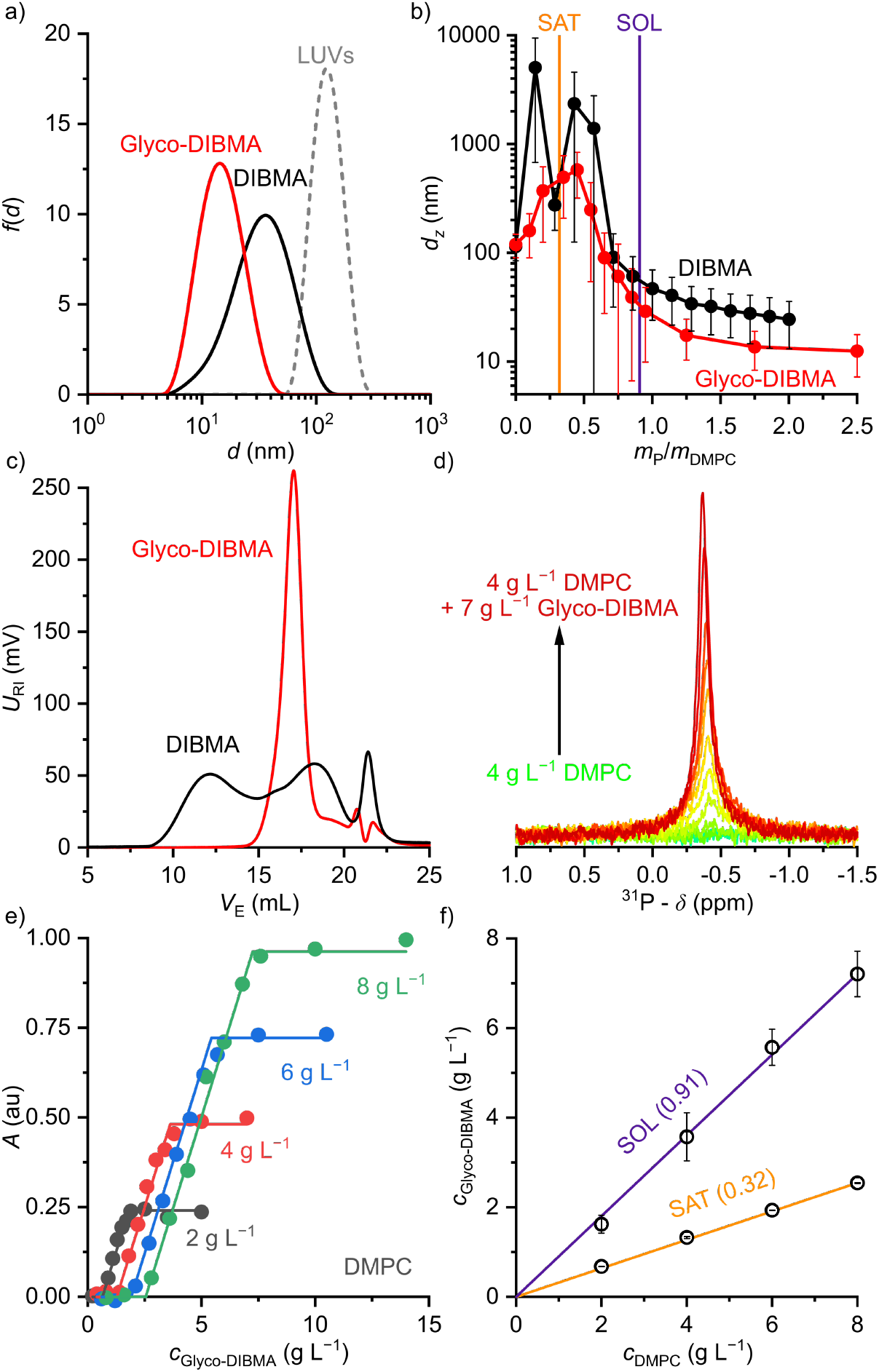
Formation of polymer-encapsulated DMPC nanodiscs by Glyco-DIBMA and DIBMA in 50 mM Tris, 200 mM NaCl, pH 7.4, at 30°C. a) Intensity-weighted particle size distributions of 3.4 g L^−1^ (5 mM) DMPC initially present in the form of LUVs before and after nanodisc formation at a polymer/lipid mass ratio of *m*_P_/*m*_L_ = 1.5. b) *z*-Average hydrodynamic diameters (*full circles*) and size distribution widths *(“error” bars*) of 3.4 g L^−1^ (5 mM) DMPC at increasing *m*_P_/*m*_L_. Vertical lines indicate saturation 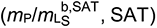 and solubilization 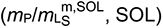 thresholds as derived by ^31^P-NMR (see panels d–f). c) SEC chromatograms showing the voltage of the refractive index detector, *U*_RI_, as a function of elution volume, *V*_E_, for DMPC nanodiscs at *m*_P_/*m*_L_ = 1.5. Minor peaks at ∼22 mL are due to NaCl and other solvent components. d) ^31^P-NMR spectra of 4 g L^−1^ (5.9 mM) DMPC at increasing concentrations of Glyco-DIBMA. e) NMR peak areas, *A*, at four different DMPC concentrations and increasing Glyco-DIBMA concentrations. Lines are from a global fit according to Eqs. 3–5 in the SI. f) Glyco-DIBMA/DMPC pseudo-phase diagram as derived from peak areas in panel e. Shown are breakpoints from local fits (*open circles*) and corresponding 95% confidence intervals (*error bars*) as well as the result of a global fit (*lines*) according to Eqs. 3–5.

We further quantified the efficiency of Glyco-DIBMA in forming nanodiscs by monitoring the emergence of a ^31^P-NMR signal upon titration of DMPC LUVs with the polymer (Figure 2d–f). As solution NMR spectroscopy is insensitive to nuclei residing in large, slowly tumbling particles such as LUVs, no peak was detectable in the absence or in the presence of low concentrations of Glyco-DIBMA (Figure 2d). Increasing *m*_P_/*m*_L_ beyond the so-called saturation (SAT) threshold resulted in the emergence of an isotropic peak, which indicated the formation of fast-tumbling Glyco-DIBMALPs. Further addition of Glyco-DIBMA caused a linear increase in the peak area until the latter reached a plateau at the so-called solubilization (SOL) boundary (Figure 2d,e). Reading the SAT and SOL breakpoints from plots of ^31^P-NMR peak area vs. Glyco-DIBMA concentrations at several DMPC concentrations (Figure 2e) furnished a pseudo-phase diagram^27,28^ (Figure 2f) characterized by SAT and SOL boundaries and associated 95% confidence intervals of, respectively, 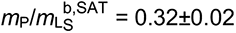 and 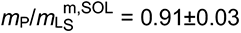 which fall in the same range as the corresponding values determined for DIBMA.^2^ Hence, nanodisc formation by Glyco-DIBMA can be rationalized in terms of a simple three-stage model,^27^ as has been shown for other amphiphilic polymers.^2,3,23,29–31^

We further explored the efficiency of Glyco-DIBMA in forming lipid-bilayer nanodiscs under different solution conditions (Figure 3a,b). To this end, we employed the unsaturated phospholipid 1-palmitoyl-2-oleoyl-*sn*-glycero-3-phosphocholine (POPC), which is more challenging to accommodate in nanodiscs than DMPC, especially for DIBMA.^3,4,23^ Indeed, incubation of POPC LUVs with DIBMA under near-physiological conditions (i.e., 50 mM Tris, 200 mM NaCl, pH 7.4, 25°C) produced large and broadly distributed nanoparticles with *d_z_ =* (85±45) nm even at a polymer/lipid mass ratio as high as *m*_P_/*m*_L_ = 6 (Figure 3a). By contrast, Glyco-DIBMA enabled the formation of narrowly distributed nanodiscs with *d_z_ =* (15±6) nm already at *m*_P_/*m*_L_ = 2.2 under otherwise identical conditions (Figure 3a). As is the case for DIBMA,^23,24^ the ionic strength of the solution substantially influenced the solubilization efficiency of Glyco-DIBMA (Figure 3b). While nanodisc formation in the absence of added salt proved difficult, increasing *c*_NaCl_ to 100 mM steadily increased the efficiency of Glyco-DIBMA to afford nanoparticles with *d_z_* ≈ (10±4) nm already at *m*_P_/*m*_L_ = 2. Although a further increase in *c*_NaCl_ beyond 100 mM slightly reduced the solubilization efficiency and increased the size of the nanoparticles, this effect was suppressed by raising the pH value to 8.3, which afforded diameters of *d_z_ =* (10±4) nm already at *m*_P_/*m*_L_ = 1 even at *c*_NaCl_ = 200 mM (Figure 3b). Moreover, Glyco-DIBMALPs were found to tolerate temperatures of up to 65°C at pH 8.3 and 200 mM NaCl (Figure SI 2). In summary, the efficiency of Glyco-DIBMA in facilitating the self-assembly of lipid-bilayer nanodiscs was greatest in the neutral to moderately alkaline pH range and at a reasonably physiological ionic strength of ∼100 mM. In contrast with DIBMA,^23^ neither acidic pH, nor high ionic strength, nor addition of divalent cations was required for optimal performance of Glyco-DIBMA, which thus proved to be a much more powerful nanodisc former under challenging conditions.

**Figure 3:**
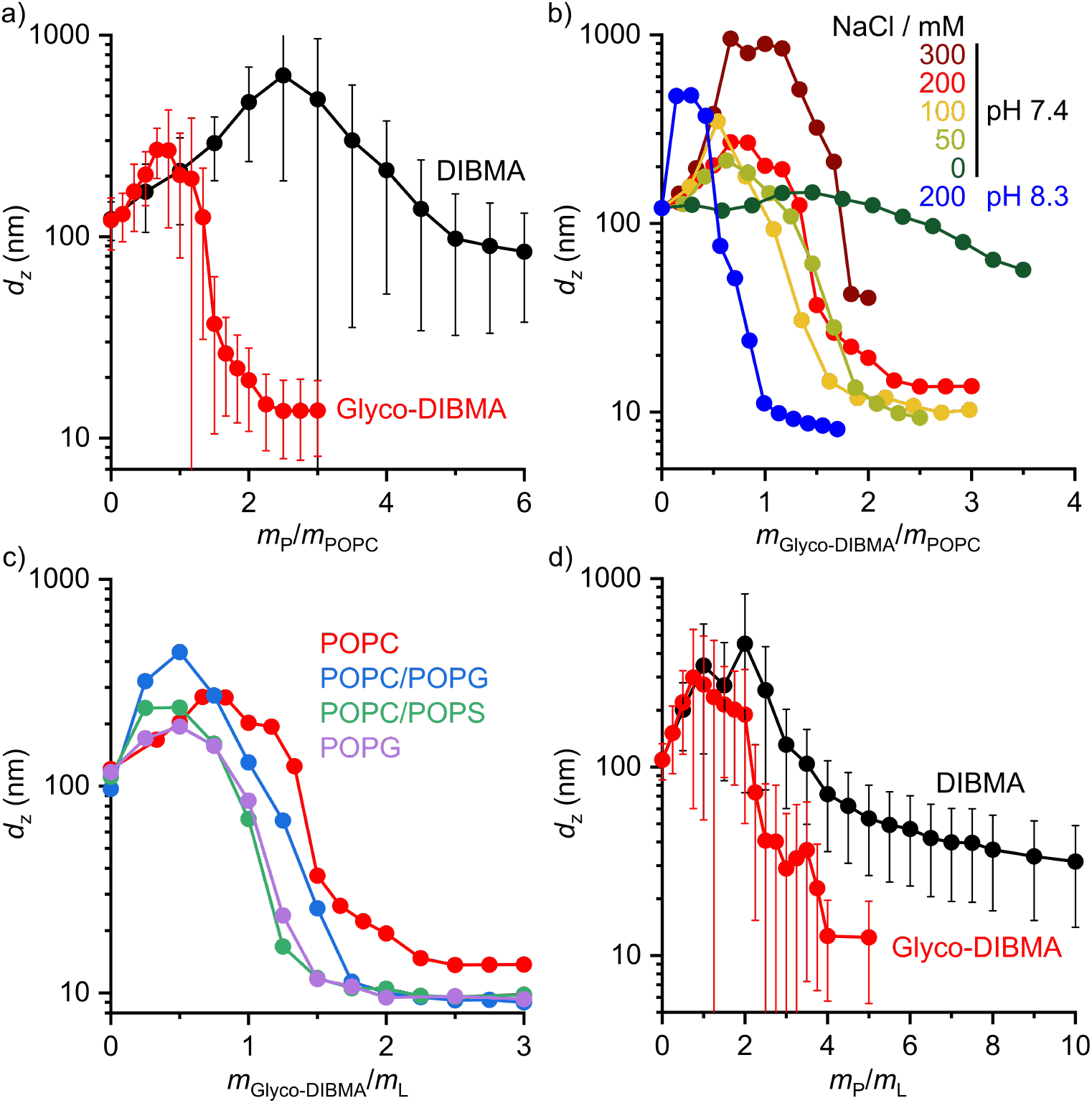
Formation of polymer-encapsulated nanodiscs by Glyco-DIBMA and DIBMA using different buffer compositions and various lipid species. a,b) *z*-Average hydrodynamic diameters of 3.8 g L^−1^ (5 mM) POPC LUVs at increasing polymer/lipid mass ratios in the presence of a) 50 mM Tris, 200 mM NaCl, pH 7.4 or b) various NaCl concentrations and pH values (for Glyco-DIBMA only). c) *z*-Average hydrodynamic diameters of LUVs harboring POPC, POPC/POPG (1:1 mol/mol), POPC/POPS (7:3 mol/mol), or POPG, each at ∼4 g L^−1^ total lipid at increasing Glyco-DIBMA/lipid mass ratios. d) *z*-Average hydrodynamic diameters of 4 g L^−1^ POPC/cholesterol (3.55:1 mol/mol) at increasing polymer/lipid mass ratios. Data shown in panels c and d were collected in 50 mM Tris, 200 mM NaCl, pH 7.4.

In addition to varying the solution conditions, we tested the performance of Glyco-DIBMA in harboring more challenging lipid mixtures in nanodiscs that more closely mimic the complex compositions of eukaryotic and prokaryotic membranes (Figure 3c,d). To this end, we incubated Glyco-DIBMA with LUVs made either from the anionic phospholipid 1-palmitoyl-2-oleoyl-*sn*-glycero-3-phospho-(1′-rac-glycerol) (POPG) or from mixtures of POPC with POPG, 1-palmitoyl-2-oleoyl-*sn*-glycero-3-phos-phoL-serine (POPS), or cholesterol. Addition of Glyco-DIBMA to all of the anionic LUVs produced nanoparticles with *d_z_* ≈ (10±4) nm already at *m*_P_/*m*_L_ = 2. By contrast, highly negatively charged DIBMA failed to solubilize any of the anionic membranes, even at ratios as high as *m*_P_/*m*_L_ = 10 (Figure SI 3). These contrasting findings again reflect the reduced charge density on Glyco-DIBMA, which results in lower Coulombic repulsion between the polymer and anionic membranes. Finally, we turned our attention to cholesterol, which does not affect the charge state of a zwitterionic phosphocholine-based lipid-bilayer membrane but increases both its hydrophobic thickness and the lateral pressure within its acyl chain region.^32,33^ Both of these effects can, in principle, be counterbalanced by an increase in the polymer density in the nanodisc rim. This, however, is limited by Coulombic repulsion among the polymer chains in the rim, especially for highly charged DIBMA. Indeed, the solubilization of LUVs containing 30 mol% cholesterol by DIBMA turned out very difficult, as nanoparticle sizes of *d_z_ =* (31±17) nm were attained only at an exceedingly high polymer/lipid mass ratio of *m*_P_/*m*_L_ = 10. By contrast, Glyco-DIBMA yielded much more narrowly sized nanoparticles with *d_z_ =* (13±7) nm already at *m*_P_/*m*_L_ = 4.

In order to assess the morphology and lipid-bilayer architecture of Glyco-DIBMALPs, we turned to negative-stain transmission electron microscopy (TEM) and differential scanning calorimetry (DSC) (Figure 4). TEM of Glyco-DIBMALPs encapsulating DMPC (Figure 4a) or POPC (Figure 4b) corroborated the formation of nanoscopic discs having diameters in excellent agreement with those determined by DLS (Figure 2a and Figure 3a). Of note, we observed numerous “rouleaux”, that is, assemblies of several nanodiscs stacking on top of one another. Although such stacks are known to be artifacts caused by the negative-staining procedure,^2,34,35^ they were helpful in that they enabled many side views of Glyco-DIBMALPs and, thus, clearly confirmed a nanodisc thickness typical of a phospholipid bilayer. With the aid of DSC, we probed the thermotropic gel-to-fluid phase transition of DMPC in Glyco-DIBMALPs at various polymer/lipid mass ratios (Figure 4c). We observed a broadening of the phase transition in Glyco-DIBMALPs as compared with polymer-free LUVs, which is indicative of nanoscopic lipid-bilayer patches displaying a reduced size of the so-called cooperative unit.^2–4^,^36^ Upon addition of Glyco-DIBMA, the melting temperature of DMPC slightly increased from *T*_m_ = 24.0°C in the absence of polymer to *T*_m_ = 24.8°C at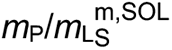, similar to what has been observed for DIBMA^2^ and SMA(2:1).^3^ Upon complete nanodisc formation at *m*_P_/*m*_L_ = 1, a value of *T*_m_ = 22°C indicated that the lipid-bilayer core encapsulated by Glyco-DIBMA was only slightly affected by the polymer. However, further addition of excess polymer caused *T*_m_ to decline to ∼13°C at *m*_P_/*m*_L_ = 4, which is similar to the effect of other polymers such as SMA(2:1).^3^

**Figure 4:**
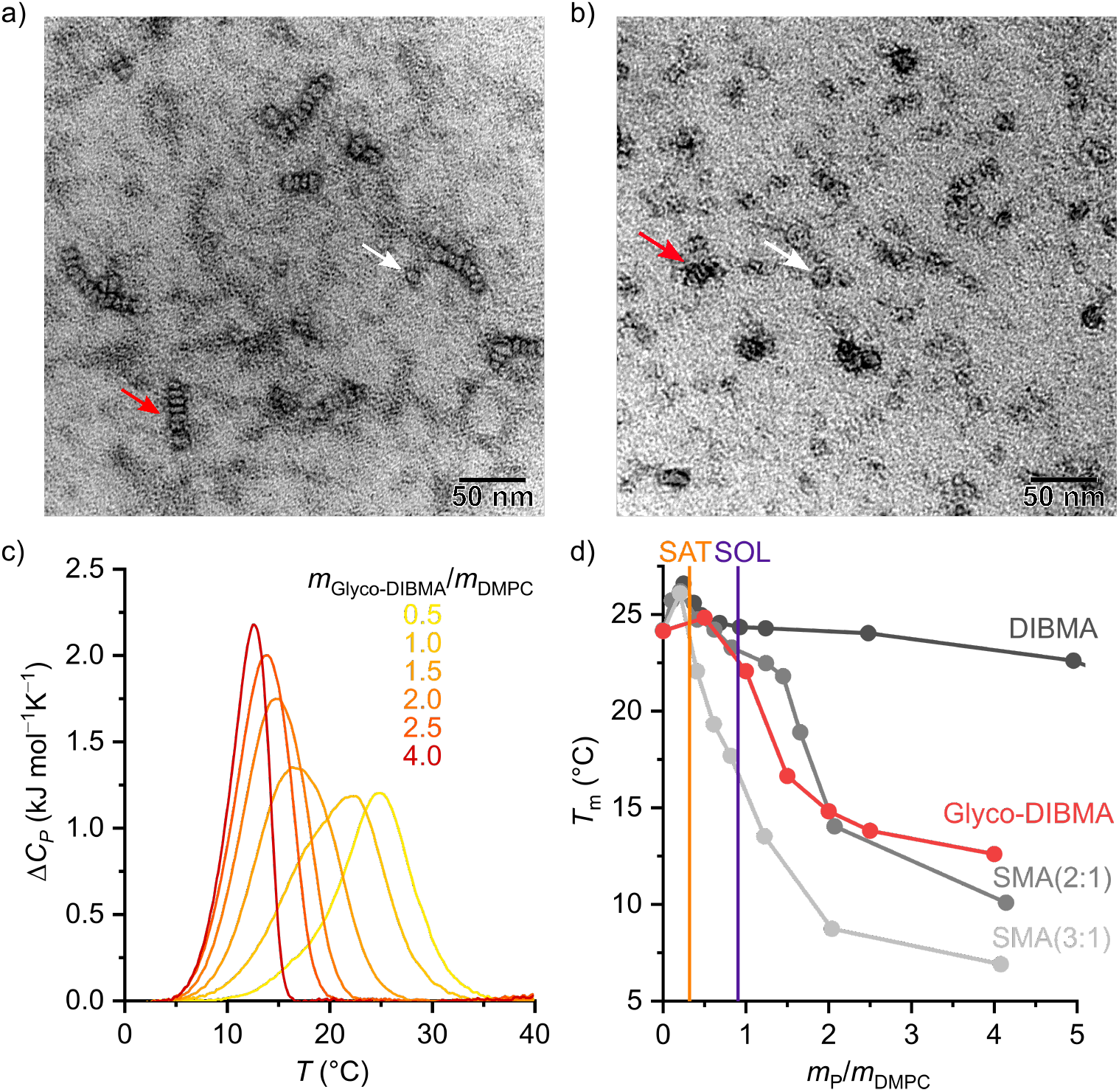
Morphology and lipid-bilayer architecture of Glyco-DIBMALPs. a,b) TEM images of negatively stained Glyco-DIBMALPs harboring a) DMPC or b) POPC at a polymer/lipid mass ratio of 1.5. Red and white arrows indicate nanodiscs in edge-on and face-on views, respectively. DSC thermograms showing excess molar isobaric heat capacities, Δ*CP*, of 3.8 g L^−1^ (5.6 mM) DMPC at increasing Glyco-DIBMA/DMPC mass ratios. d) Gel-to-fluid phase transition temperatures, *T*_m_, of DMPC as functions of polymer/DMPC mass ratio for Glyco-DIBMA (this work), DIBMA,^2^ SMA(2:1),^3^ and SMA(3:1).^2^ Vertical lines indicate saturation 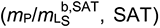 and solubilization 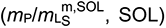 thresholds for Glyco-DIBMA as obtained from ^31^P-NMR (see Figure 2d–f).

The stronger impact on *T*_m_ of Glyco-DIBMA and the aromatic SMA copolymers as compared with DIBMA is readily explained by the smaller sizes of Glyco-DIBMALPs (Figure 2b) and SMALPs,^3,30^ respectively, which lead to a more pronounced exposure of encapsulated phospholipid molecules to the polymer rim. Conversely, these observations suggest that the chemical nature of the polymer moieties in direct contact with the lipid core (i.e., diisobutylene or styrene) plays only a minor role in determining a polymer’s effect on the thermotropic phase transition of the encapsulated lipid bilayer. Furthermore, it should be kept in mind that virtually all Glyco-DIBMA chains but only a fraction of DIBMA chains participate in nanodisc formation (Figure 2c), so that the effective polymer/lipid ratios in DIBMALPs are lower than the nominal values. Together, TEM and DSC confirmed that the solubilization of vesicular membranes by Glyco-DIBMA resulted in the formation of lipid-bilayer nanodiscs, as desired, rather than mixed micellar structures, even at high mass ratios.

It has been shown that polymer-encapsulated nanodiscs can exchange their contents with each other^4,24,37,38^ or with other lipidic systems such as vesicles,^30^ lipid monolayers,^39^ planar lipid bilayers,^40^ and cubic phases.^11^ This fast exchange of contents is enabled by the flexible, soft nature of the polymer rim, which itself exchanges rapidly among nanodiscs.^41^ Importantly, DIBMALPs and SMALPs therefore represent equilibrium nanocomposites rather than kinetically trapped lipid assemblies such as LUVs^42^ or nanodiscs bounded by membrane scaffold proteins (MSPs).^43^ Under typical *in vitro* conditions (i.e., millimolar lipid concentrations), lipid exchange occurs predominantly through nanodisc collisions^4,24,37,38^ and, consequently, slows down with increasing charge density in the polymer rim.^4,24,38^ Following this rationale, we expected lipid exchange among Glyco-DIBMALPs to be faster than among nanodiscs suffering from high charge densities such as DIBMALPs. To test this prediction, we quantified the kinetics and unraveled the mechanisms of lipid transfer by time-resolved Förster resonance energy transfer (TR-FRET). To this end, we produced fluorescently labeled nanodiscs consisting of a DMPC matrix hosting 1 mol% of each *N*-(7-nitrobenz-2-oxa-1,3-diazol-4-yl)-1,2-dihexadecanoyl-*sn*-glycero-3-phosphoethanolamine (NBD-PE) and *N*-(lissamine rhodamine B sulfonyl)-1,2-dihexadecanoyl-*sn*-glycero-3-phosphoethanolamine (Rh-PE), which served as donor and acceptor fluorophores, respectively. When co-localized within the same nanodisc, NBD-PE and Rh-PE form an efficient FRET pair, but redistribution of the fluorescent lipids in a larger lipid pool will dequench the donor and cause an increase in NBD-PE fluorescence.

Indeed, mixing labeled and unlabeled Glyco-DIBMALPs at various lipid concentrations resulted in a strong increase in the emission intensity of the donor dye NBD-PE in a time-and concentration-dependent manner (Figure 5a). Local fits (Eq. 9 in the SI) yielded transfer rate constants, *k*_obs_, similar to those retrieved from a global fit (Eq. 10) across all lipid concentrations tested, showing a linear increase in *k*_obs_ except at the lowest lipid concentration (Figure 5b). These are the hallmarks of two independent lipid-exchange mechanisms, with diffusional lipid transfer playing a significant role only at low lipid concentrations <1 mM and collisional lipid transfer dominating at higher concentrations (Figure 5c). We found the diffusional and collisional transfer rate constants and their associated 95% confidence intervals to amount to, respectively, *k*_dif_ = (0.048±0.002) s^−1^ and *k*_col_ = (43.1±0.5) M^−1^ s^−1^. Thus, the second-order collisional lipid transfer among Glyco-DIBMALPs is ∼30-fold faster than among DIBMALPs,24 which again can be ascribed to a substantial reduction in the charge density on the polymer chain upon glycosylation. In agreement with this interpretation, the overall lipid-exchange rate constant, *k*_obs_, among nanodiscs encapsulated by Glyco-DIBMA falls between those bounded by SMA(2:1) and SMA(3:1) (Figure 5d), which have considerably lower charge densities than DIBMA.

**Figure 5:**
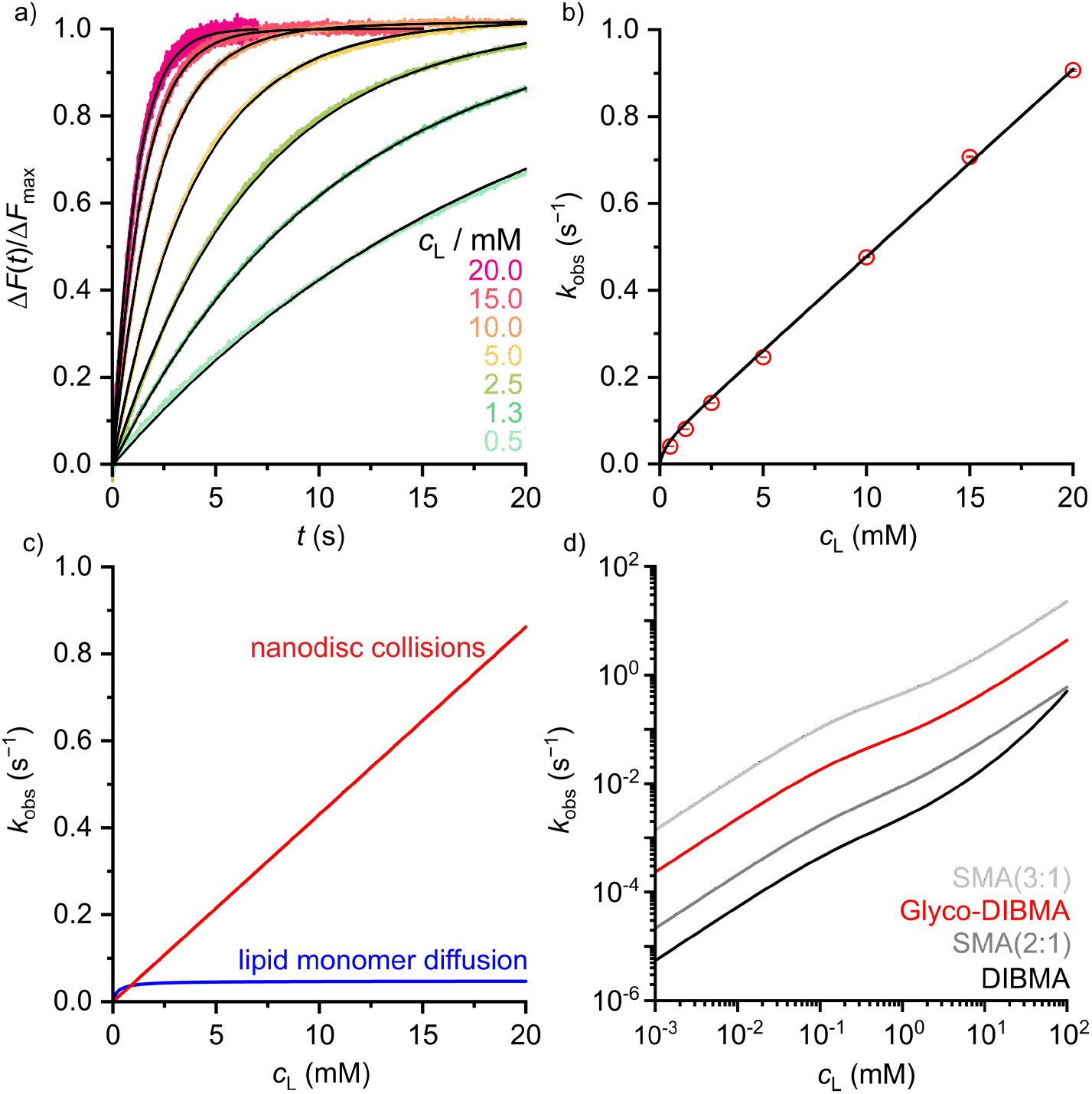
Lipid transfer among fluorescently labeled and unlabeled DMPC nanodiscs bounded by Glyco-DIBMA. a) Normalized fluorescence emission intensity of NBD-PE, Δ*F*(*t*)/Δ*F*_max_, upon rapid mixing of labeled and unlabeled nanodiscs at various lipid concentrations. Shown are experimental data points (*colored dots*) and a global fit (*black lines*) based on Eq. 10. b) Observed lipid-transfer rate constants, *k*_obs_, as functions of lipid concentration, *c*_L_. Shown are local fits (*empty circles*) and a global fit (*black line*) based on Eqs. 9 and 10, respectively. Error bars (within circles) correspond to 95% confidence intervals from local fits. c) Contributions of diffusional and collisional lipid-transfer mechanisms to *k*_obs_ as derived from Eqs. 6 and 7, respectively. d) *k*_obs_ values as functions of *c*L as determined for nanodiscs bounded by Glyco-DIBMA (this work), DIBMA,^24^ SMA(2:1),^38^ or SMA(3:1).^37^

After testing Glyco-DIBMA with respect to the encapsulation of well-defined model lipid bilayers within nanodiscs, we explored its usefulness for extracting and accommodating integral membrane proteins from complex, cellular membranes in nanodiscs. For this purpose, we first exposed *Escherichia coli* membranes to increasing concentrations of Glyco-DIBMA and quantified the total amounts of extracted membrane proteins by sodium dodecyl sulfate polyacrylamide gel electrophoresis (SDS-PAGE) (Figure 6). Across all polymer concentrations, Glyco-DIBMALPs were able to encapsulate a large variety of membrane proteins from both the inner and the outer bacterial membranes,^44,45^ revealing an SDS-PAGE band pattern similar to that found for DIBMA (Figure 6a) and other, aromatic polymers.^2,3^ At low polymer concentrations of 0.3% (*w*/*v*), both aliphatic copolymers performed roughly on par, but Glyco-DIBMA outperformed DIBMA by more than 2.5-fold at more commonly used polymer concentrations of 0.5% and 1.0% (*w*/*v*) (Figure 6b). As *E. coli* membranes are highly anionic,^46^ nanodisc-forming polymers need to overcome strong Coulombic repulsion in order to interact with and fragment such membranes. This explains why, for optimal performance, partial charge compensation by divalent cations is required for highly charged DIBMA^4^ but not for Glyco-DIBMA, which revealed a much higher protein-extraction efficiency under mild, near-physiological conditions.

**Figure 6:**
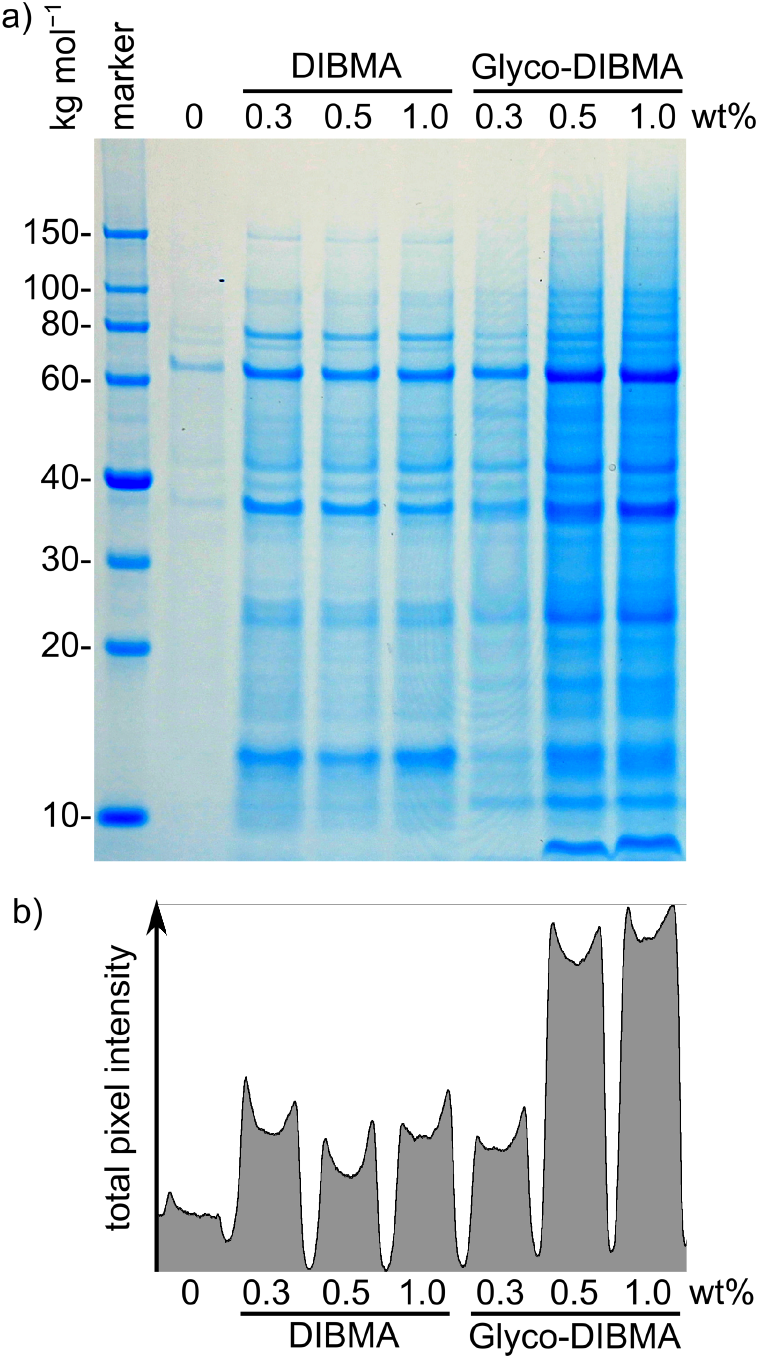
Extraction of membrane proteins from *E. coli* cells into polymer-encapsulated nanodiscs mediated by Glyco-DIBMA and DIBMA. Shown are a) a Coomassie-stained gel after SDS-PAGE of polymer-solubilized fractions and b) a projection of the total pixel intensity across all lanes in the gel. Cell debris and water-soluble proteins were removed by serial ultracentrifugation, and samples were gently agitated overnight at 23°C in the presence of Glyco-DIBMA or DIBMA at a constant *E. coli* membrane concentration of 5% (*w*/*v*). Prior to electrophoresis, unsolubilized material and polymer were removed by ultracentrifugation and organic solvent extraction, respectively. A control without polymer was produced under otherwise identical conditions.

Next, we sought to demonstrate that Glyco-DIBMA not only extracts membrane proteins from cellular membranes to accommodate them in nanodiscs but can also be used to purify and chemically manipulate a particular target protein directly within Glyco-DIBMALPs (Figure 7). Hence, we extracted a His_6_-tagged variant of the voltage-gated K^+^ channel KvAP^47^ from *E. coli* membranes and purified it by Co^2+^ affinity chromatography and SEC. This two-step purification resulted in a single strong, band at the expected molar mass of ∼25 kg mol^−1^ on SDS-PAGE, both in a Coomassie stain (Figure 7a) and in a Western blot relying on an antibody directed against the His_6_-tag (Figure 7b). Mass spectrometry (MS) confirmed that this band corresponded to KvAP. Of note, SEC (Figure 7c) allowed neat separation of KvAP from SlyD (Figure 7a, fraction 5), a histidine-rich protein that is a common contaminant in the affinity purification of His-tagged proteins,^48,49^ to afford excellent purity of KvAP in Glyco-DIBMALPs (Figure 7a,b). Fraction 3 of the SEC run was thus concentrated and subjected to further analysis by DLS (Figure 7d) and TEM (Figure 7e) to confirm that it contained discoidal nanoparticles with an average diameter of ∼15 nm.

**Figure 7:**
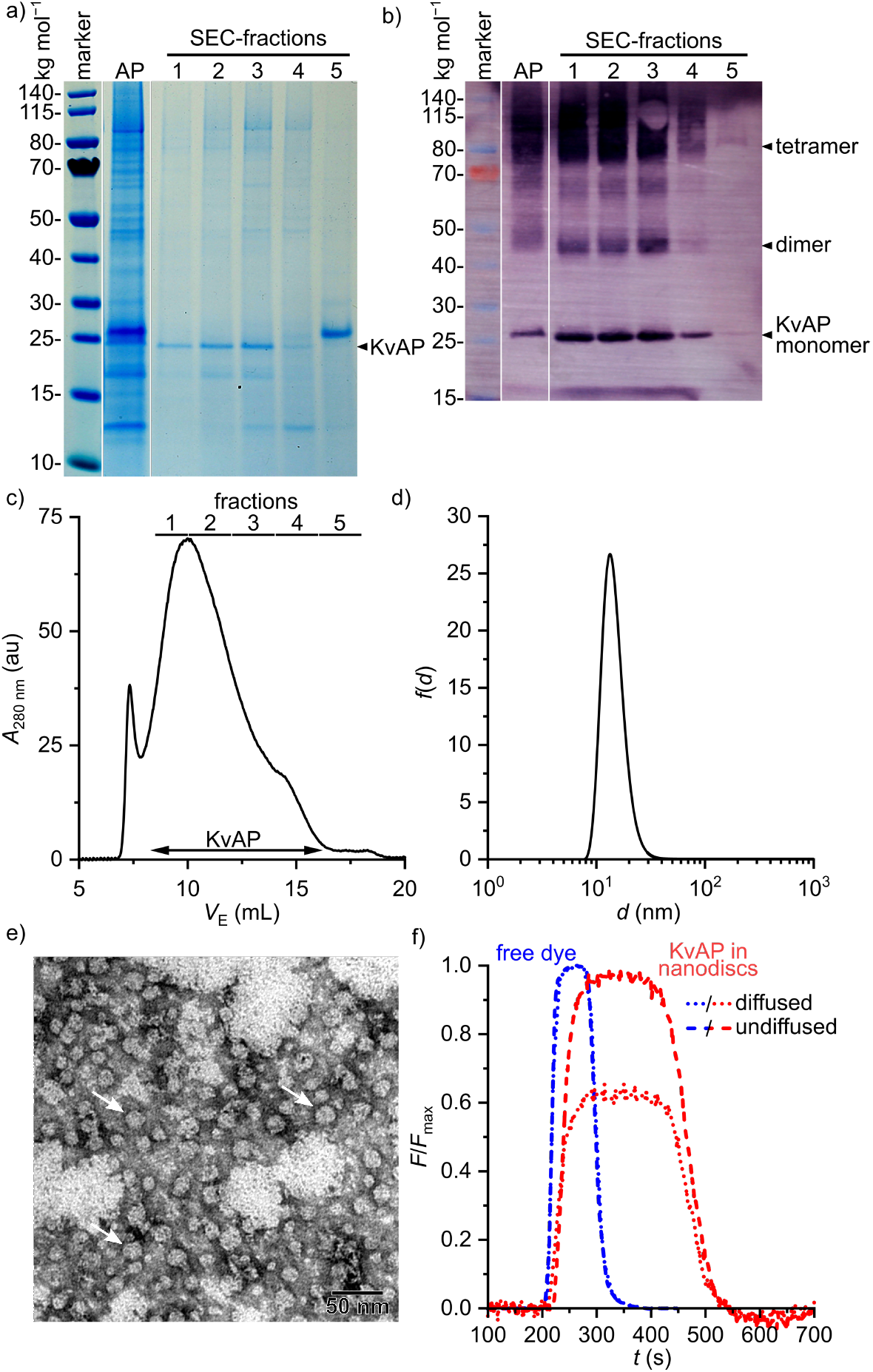
Extraction and purification of the voltage-gated K^+^ channel KvAP embedded in Glyco-DIBMA nanodiscs. a,b) SDS-PAGE of KvAP extracted and purified with 5% (*w*/*v*) Glyco-DIBMA as visualized by a) Coomassie stain and b) Western blot. Shown are samples after His_6_-Tag affinity purification and concentration (*AP*) and after SEC as indicated in panel c. c) Representative SEC elution profile of the concentrated sample (*AP*) shown in panels a and b on a Superose 6 Increase column. Shown is the absorption at 280 nm as a function of elution volume,*V*_E_. Number-weighted particle size distribution of Glyco-DIBMALPs harboring purified KvAP, as concentrated from SEC fraction 3 shown in panels a–c. e) TEM image of KvAP-containing Glyco-DIBMALPs (same sample as shown in panel d). White arrows indicate nanodiscs in face-on views. f) Normalized fluorescence emission intensity at ∼520 nm of labeled KvAP (from SEC fraction 3) and unconjugated ATTO-488 maleimide after injection into a microfluidic laminar flow chamber and separation into two detection channels corresponding to diffused (*dotted lines*) and undiffused fluorophores (*dashed lines*).

As a final proof of concept, microfluidic diffusional sizing (MDS)^50^ was used to show that protein-containing Glyco-DIBMALPs are amenable to protein-conjugation techniques and further *in vitro* scrutiny. In contrast with DLS and TEM, MDS selectively reports on the hydrodynamic size of fluorescently labeled nanoparticles only. Hence, we labeled affinity-purified KvAP embedded in Glyco-DIBMALPs by conjugation of the thiol-reactive fluorophore ATTO-488 maleimide to the protein’s sole cysteine residue at position 243. After removal of unconjugated dye by SEC, labeled KvAP incorporated in Glyco-DIBMALPs was subjected to MDS and compared with the free, unconjugated dye. The MDS raw data revealed that the fraction of diffused KvAP was much smaller than that of the free dye (Figure 7f), as expected from the large increase in size of the fluorescent species upon conjugation to a nanodisc-embedded membrane protein. Quantitative analysis of the MDS data furnished a diffusion coefficient of 3.8 μm^2^ s^−1^ and a corresponding hydrodynamic diameter of 11 nm for KvAP nanodiscs, whereas free ATTO-488 maleimide displayed a diffusion coefficient of 300 μm^2^ s^−1^ and a hydrodynamic diameter of 1.5 nm. These values are in excellent agreement with structural considerations and, thus, demonstrate that nanodiscs encapsulsated by Glyco-DIBMA are compatible with chemical protein-conjugation techniques, state-of-the-art microfluidic methods, and sensitive fluorescence detection without ever removing the target protein from its native-like lipid-bilayer background.

## Conclusions

We have introduced Glyco-DIBMA as a more hydrophobic, less charged, and more powerful nanodisc-forming polymer than established aliphatic membrane-solubilizing copolymers. This new, bioinspired glycopolymer fragments and accommodates model and cellular membranes with substantially higher efficiency—in particular, under near-physiological conditions—to form smaller and more narrowly distributed lipid-bilayer nanodiscs. Glyco-DIBMALPs harbor a native-like, dynamic lipid-bilayer core and are stable across a wide range of buffer compositions and temperatures. Glyco-DIBMALPs are able to accommodates within their lipid-bilayer core a broad range of membrane proteins, which thus become amenable to chromatographic purification as well as further *in vitro* manipulation and analysis while remaining embedded in a nanoscale membrane patch at all times.

## Acknowledgements

We thank Dr. Harald Kelm, Lisa Hamsch, Christiane Müller, Ann-Cathrin Schlapp, Dr. Frederik Sommer, and Kai Zwara (all TUK) for excellent technical assistance and Prof. Peter Pohl (University of Linz) for providing the *E. coli* strain used for producing KvAP. This work was supported by the Carl Zeiss Foundation through the Centre for Lipidomics (CZSLip). M.T.A. acknowledges the Deutsche Akademische Austauschdienst (DAAD) for a Ph.D. scholarship.

## Notes

### Competing Interest Statement

Dr. Cenek Kolar is the founder and owner of Glycon Biochemicals GmbH, which will offer Glyco-DIBMA for sale after publication of this manuscript. Marie Rasche is an employee of Glycon Biochemicals GmbH.

