## Supporting Information for "A Bioinspired Glycopolymer for Capturing Membrane Proteins in Native-Like Lipid-Bilayer Nanodiscs"

### 1. Experimental section

#### Materials

All chemicals were purchased in the highest available purity. DIBMA (trade name ACUSOL 460ND, $M_w$ = 10 kg mol$^{-1}$) was purchased from Brenntag (Essen, Germany) and meglumine was purchased from Merck (Darmstadt, Germany). Acetic anhydride and toluene were purchased from OQEMA (Korschenbroich, Germany). DMPC and POPC were kind gifts from Lipoid (Ludwigshafen, Germany). POPE and POPS were purchased from Avanti Polar Lipids (Alabaster, USA), POPG was from Genzyme (Cambridge, USA), and cholesterol was from Sigma–Aldrich (Steinheim, Germany). NBD-PE and Rh-PE were purchased from Fisher Scientific (Schwerte, Germany) and Biotium (Fremont, USA), respectively. Atto-488 maleimide, isopropyl-β-D-thio¬galacto¬pyranoside (IPTG), tris(2-carboxyethyl)phosphine (TCEP), dithiothreitol (DTT), anti-mouse IgG, 85% (*w*/*v*) $H_3PO_4$ in $D_2O$, $CDCl_3$, and $CD_3COD$ were purchased from Sigma-Aldrich. Albumin fraction V, Coomassie Brilliant Blue G250, nitrotetrazolium blue chloride (NBT), 5-bromo-4-chloro-3-indolyl phosphate (BCIP), 4-(2-hydroxyethyl)-1-piperazineethanesulfonic acid (HEPES), tris(hydroxyl-methyl)aminomethane (TRIS), ethylenediaminetetraacetic acid (EDTA), and sodium dodecyl sulfate (SDS) were from Carl Roth (Karlsruhe, Germany). 6x-His-tag monoclonal antibody was from Invitrogen (Rockford, USA), and Tween-20 from Serva (Heidelberg, Germany). Pure Na, benzonase nuclease, and imidazole were from Merck. $D_2O$ was purchased from Deutero (Kastellaun, Germany), and NaCl, KCl, $CHCl_3$, and $CH_3OH$ were from VWR (Darmstadt, Germany).

#### Synthesis of Glyco-DIBMA

Conversion of DIBMA acid to anhydride: A solution of DIBMA acid (22.8 g) in acetic anhydride (41 mL) was heated in a 500-mL round-bottomed flask at 50°C until it became clear. After addition of toluene (40 mL), the solution was stirred at 70°C for 50 min before the solvent was removed *in vacuo*. The resulting product was distilled twice with toluene (40 mL) and dried in a vacuum oven at 36°C for 48 h. The yield of dry colorless product was 19 g (90%). Amidation of DIBMA anhydride: Sodium (1.2 g) dissolved in methanol (50 mL) and *N*-methyl-D-glucamine (9.8 g) were added to a solution of DIBMA anhydride (10.5 g) in methanol (100 mL). The solution was stirred and refluxed at 65°C for 8 h. Ethanol (100 mL) was added to the reaction mixture, and the solvent was removed *in vacuo*. The solid product obtained was dried in a vacuum oven at 36°C for 48 h. The yield of dry colorless product was 21 g (98%).

#### Preparation of polymer stock solutions

3.5 mL of 15–20% (*w*/*v*) polymer solution was produced by dissolving Glyco-DIBMA or DIBMA powder in ultrapure water at 50°C. The resulting polymer solutions were dialyzed against 800 mL buffer (50 mM Tris, 200 mM NaCl, pH 7.4 or 8.3) using 5-mL QuixSep dialyzer capsules (Carl Roth) using Spectra/Por 3 dialysis membranes with a nominal molecular weight cut-off of 3.5 kg/mol (Spectrum Laboratories, Rancho Dominguez, USA). Dialysis was performed for 24 h at room temperature with buffer exchange after 16 h. Dialyzed polymer solutions were sterile-filtered using 220-nm poly(vinylidene fluoride) syringe filters (Carl Roth). Glyco-DIBMA concentrations were determined by refractometry on an Abbemat 500 (Anton Paar, Graz, Austria) using a specific refractive index increment of $dn/d\rho$ = 0.151 L kg$^{-1}$ at 589 nm or a specific extinction coefficient of $\varepsilon$ = 1.62 L (g cm)$^{-1}$ at 220 nm. DIBMA concentrations were determined by using $dn/d\rho$ = 0.14 L kg$^{-1}$ at 589 nm or $\varepsilon$ = 3.49 L (g cm)$^{-1}$ at 220 nm. Both refractive and absorption measurements were performed at 20°C.

#### Preparation of LUVs

POPC, POPG, POPS, and cholesterol were separately dissolved in $CHCl_3$ and then mixed at a molar ratio of 1:1 (POPC/POPG), 7:3 (POPC/POPS), and 3.55:1 (POPC/cholesterol). A thin lipid film was obtained after solvent removal using a continuous nitrogen stream. Trace amounts of solvent were removed under high vacuum in a desiccator for at least 16 h. Lipid films were resuspended in buffer (50 mM Tris, 200 mM NaCl, pH 7.4) by vortexing at 35°C, which was followed by at least six freeze–thaw cycles. For LUVs consisting of a single lipid species, lipid powder was directly suspended in 50 mM Tris, 0–200 mM NaCl, pH 7.4 or pH 8.3 to reach $c_L$ = 20–50 g L$^{-1}$. All lipid suspensions were heated to 40°C and vortexed for 2 min prior 21–31-fold extrusion through two stacked polycarbonate membranes with a nominal pore diameter of 100 nm. Extrusion of DMPC and POPC/cholesterol mixtures was performed at 35°C using a block-heated Mini-Extruder (Avanti, Alabama, USA), whereas all other lipids and lipid mixtures were extruded at room temperature using a LiposoFast extruder (Avestin, Ottawa, Canada). DLS (see below) confirmed unimodal particle size distributions, yielding hydrodynamic LUV diameters of ~140 nm for DMPC, ~160 nm for POPC, ~120 nm for POPG, ~110 nm for POPC/cholesterol, ~100 nm for POPC/POPG, and ~110 nm for POPC/POPS.

#### Solubilization efficiency probed by DLS

LUVs were mixed with Glyco-DIBMA or DIBMA to yield final concentrations of 3.3–3.9 g L$^{-1}$ (5 mM) of total lipid and either 0–25 g L$^{-1}$ Glyco-DIBMA or 0–100 g L$^{-1}$ DIBMA. Samples were incubated under shaking at 500 rpm for 16 h at 25°C for POPC, POPG, and mixtures of POPC/POPG or POPC/POPS, at 35°C for DMPC, and at 37°C for mixtures of

POPC/cholesterol. For POPC, we repeated solubilization experiments at 0–300 mM NaCl at pH 7.4 as well as at 200 mM NaCl and pH 8.3. Particle sizes and size distributions were monitored by DLS (see below).

**Solubilization efficiency probed by $^{31}$P-NMR spectroscopy**

Samples containing 2, 4, 6, and 8 g $L^{-1}$ DMPC and 0–14 g $L^{-1}$ Glyco-DIBMA were incubated for 16 h at 35°C. 10% $D_2O$ (*v*/*v*) was included in each sample to provide a lock signal. $^{31}$P-NMR measurements were carried out at 30°C on an Avance 600 spectrometer (Bruker Biospin, Rheinstetten, Germany) operating at a $^{31}$P-resonance frequency of 242.9 MHz using a 5-mm broadband inverse probe. 128 scans were acquired with an inverse-gated decoupling sequence using an acquisition time of 2.2 s, a sweep width of 7310 Hz, and a relaxation delay of 6 s. Data were multiplied by an exponential function with a line-broadening factor of 1.0 Hz before Fourier transformation. Chemical shifts were referenced to 85% (*w*/*v*) $H_3PO_4$ in $D_2O$ as external standard at 0 ppm. Peaks were integrated using the software Bruker Topspin 4.0.9.

**Preparation of nanodiscs for TR-FRET**

For fluorescently labeled nanodiscs, DMPC, NBD-PE, and Rh-PE powders were separately dissolved in $CHCl_3$ and then mixed at a molar ratio of 98:1:1. A thin lipid film was obtained after solvent removal using a continuous nitrogen stream. Trace amounts of solvent were removed under high vacuum in a desiccator for at least 16 h, and the lipid film was resuspended in buffer. For unlabeled nanodiscs, DMPC powder was directly suspended in buffer. Both labeled and unlabeled lipid stocks were vortexed at 35°C, which was followed by at least six freeze–thaw cycles. Lipid suspensions were mixed with Glyco-DIBMA at final concentrations of 3.4 g $L^{-1}$ (5 mM) or 27.1 g $L^{-1}$ (40 mM) total lipid and equal mass concentrations of Glyco-DIBMA for labeled and unlabeled nanodiscs, respectively. Both mixtures were incubated at 35°C on a thermoshaker at 500 rpm for 16 h. For both labeled and unlabeled nanodiscs, unimodal size distributions were confirmed by DLS (see below), yielding *z*-average hydrodynamic diameters, $d_z$, and associated size-distribution widths of $d_z$ = (22±5) nm. Finally, labeled nanodiscs were diluted to a total lipid concentration of 0.3 g $L^{-1}$ (0.5 mM), whereas unlabeled nanodiscs were diluted to lipid concentrations of 20.3, 13.6, 6.8, 3.4, 1.7, and 0.7 g $L^{-1}$ (30, 20, 10, 5, 2.5, and 1 mM, respectively).

**TR-FRET**

We followed time-dependent donor dequenching on an SF.3 stopped-flow apparatus equipped with a (470±10) nm light-emitting diode, whose output was set to 10–20 mA, further attenuated by an OD 2 filter to avoid photobleaching. Fluorescence emission was blocked below 513 nm and above 543 nm with a TechSpec OD 6 band-pass filter (Edmund Optics, Karlsruhe, Germany) and was monitored with a photomultiplier mounted at a right angle. Drive syringes, tubes, and the quartz glass cell were thermostatted at 30°C at all times. Samples were equilibrated for at least 10 min prior to each measurement. Then, 75-µL aliquots of fluorescently labeled nanodiscs at a total lipid concentration of 0.3 g $L^{-1}$ (0.5 mM) were mixed rapidly with equal volumes of unlabeled nanodiscs at lipid concentrations of 27.1, 20.3, 13.6, 6.8, 3.4, 1.7, and 0.7 g $L^{-1}$ (40, 30, 20, 10, 5, 2.5, and 1 mM, respectively). Time-dependent emission from NBD-PE was recorded 5 times with 10,000 data points per shot. Fluorescence transients thus obtained were averaged and analyzed by nonlinear least-squares fitting.[1]

**DLS**

DLS measurements were carried out on a Zetasizer Nano S90 (Malvern Panalytical, Malvern, UK) equipped with a He–Ne laser emitting at 633 nm. Samples were measured in a 3 mm × 3 mm quartz glass cuvette (Hellma Analytics, Müllheim, Germany) with automated laser attenuation. For LUVs containing POPC or mixtures of POPC/POPG and POPC/POPS, the cuvette was thermostatted at 25°C, for DMPC at 30°C, and for mixtures of POPC and cholesterol at 37°C. To test the colloidal stability of Glyco-DIBMALPs harboring POPC as a function of temperature, we repeated DLS measurements at 5–75°C. Prior to measurements, the samples were equilibrated for at least 15 min at each temperature. Autocorrelation functions were fitted using a non-negatively constrained least-squares function[2] to yield intensity-weighted particle size distributions and by cumulant analysis[3] to obtain $d_z$ values and associated polydispersity indices (PDIs). Distribution widths of $d_z$, $\sigma$, were calculated as $\sigma = \sqrt{\text{PDI}}\, d_z$.

**DSC**

Samples containing 3.8 g $L^{-1}$ (5.6 mM) DMPC and 0.3–15.2 g $L^{-1}$ Glyco-DIBMA were incubated at 35°C for 16 h. Sample and reference cells were filled with buffer and were repeatedly heated and cooled at a rate of 30°C $h^{-1}$ before the buffer in the sample cell was replaced with sample. Apart from the first upscan, successive heating and cooling scans, which were also performed at a rate of 30°C $h^{-1}$, overlaid very closely. Data were averaged, blank-subtracted, and normalized against the molar amount of DMPC in the sample using the software MicroCal Origin 7.0 (OriginLab, Northampton, USA). The melting temperature, $T_m$, was taken as the temperature at which the excess molar isobaric heat capacity, $\Delta C_p$, reached a maximum.

**Analytical SEC**

Analytical SEC was performed on an OmniSEC system (Malvern Panalytical, Malvern, UK) equipped with a Superose 6 Increase 10/300 GL column (Cytiva, Freiburg, Germany). Samples, the column, and the detector were thermostatted at 23°C or 30°C. The column was equilibrated with buffer (50 mM Tris, 200 mM NaCl, pH 7.4) at a steady flow rate of 0.5 mL $min^{-1}$ for at least three column volumes (i.e., >75 mL) before a 50-μL aliquot of the sample was injected.

**Protein extraction from cellular membranes**

*E. coli* BL21(DE3) cells were transformed with an empty pET-24 vector and selected by kanamycin resistance. After incubation in lysogeny broth (LB) overnight at 37°C under permanent agitation, cells were harvested by centrifugation at 7000 *g* for 15 min and washed with saline (154 mM NaCl). Cell pellets were resuspended in ice-cold 100 mM $Na_2CO_3$ (10 mL $g^{-1}$ cell pellet) and ultrasonicated twice with two 10-min cycles consisting of alternating 1-s pulses and 1-s breaks at an amplitude of 40% with a 10-min break between the two cycles using a Sonopuls MS73 (Bandelin, Berlin, Germany). Cell debris and unbroken cells were removed by centrifugation at 3000 *g* and 4°C for 30 min. The supernatant was subjected to ultracentrifugation for 1 h at 100,000 *g* and 4°C and washed with buffer (50 mM Tris, 200 mM NaCl, 2 mM EDTA, pH 7.4). Membrane pellets were resuspended in the same buffer to a final concentration of 50 g $L^{-1}$ wet mass and treated with 0–1% (*w*/*v*) Glyco-DIBMA or DIBMA. Samples were incubated at 20°C for 16 h with shaking at 500 rpm before ultracentrifugation for 1 h at 100,000 *g* and 4°C. The supernatant containing solubilized membrane proteins was analyzed by SDS-PAGE. To avoid band smearing by polymers,[4] solubilized fractions were precipitated with $CH_3OH/CHCl_3/H_2O$ at a mixing ratio of 4:1:3 (*v*/*v*/*v*).[5] Briefly, to a 200-μL aliquot of ice-cold sample, we successively added 800 μL $CH_3OH$, 200 μL $CHCl_3$, and 600 μL water with thorough vortexing after each addition. The mixture was centrifuged for 2 min at 14,000 *g* and 4°C. The upper, aqueous layer was removed, and 800 μL $CH_3OH$ was added before the sample was vortexed again. Precipitated proteins were pelleted by centrifugation for 1 min at 5000 *g* and another 5 min at 20,000 *g*, both at 4°C. $CH_3OH$ was carefully removed using a pipette without disturbing the pellet. Residual organic solvent was evaporated in a Jouan RC1010 vacuum concentrator (Fisher Scientific). The dried pellet was resuspended in 100 μL 2-fold concentrated SDS loading buffer (106 mM Tris-HCl, 141 mM Tris, 2% (*w*/*v*) SDS, 10% (*w*/*v*) glycerol, 0.51 mM EDTA, 0.22 mM Coomassie Brilliant Blue G250, 0.175 mM Phenol Red, 25 mM DTT, pH 8.5), boiled at 95°C for 10 min at 1000 rpm, and subjected to SDS-PAGE (see below).

**Production, extraction, and purification of KvAP**

3 L of LB medium containing 100 mg $L^{-1}$ ampicillin was inoculated with 40 mL of an overnight preculture of *E. coli* BL21(DE3) containing the plasmid pQE60 harboring the cDNA of KvAP with a C-terminal hexa-histidine ($His_6$) tag and was incubated at 37°C under constant agitation.[6] When the cultures reached $OD_{600}$ = 0.8–1.0, expression was induced by the addition of 0.4 mM IPTG supplemented with 10 mM $BaCl_2$ for 4 h at 37°C. Cells were harvested by centrifugation (7000 *g*, 15 min at 4°C) and washed with 25 mM Tris, 100 mM KCl, 1 mM $MgCl_2$, at pH 8.0. 7.1 g of the cell pellet was resuspended in the same buffer containing EDTA-free protease inhibitors and benzonase (1:3 *w*/*v*). Cell disruption was performed by ultrasonication using a Sonopuls MS73 (Bandelin) performing 5 cycles of 6 min each with 5-s pulses interrupted by 10-s breaks at an amplitude of 40%. The cell lysate was centrifuged for 10 min at 3000 *g* and 4°C, and the supernatant was ultracentrifuged for 90 min at 100,000 *g* and 4°C. The resulting membrane fraction was resuspended to 60 g $L^{-1}$ in 25 mM Tris, 100 mM KCl, 1 mM $MgCl_2$, and 5% (*w*/*v*) Glyco-DIBMA at pH 8.0. Solubilization was performed over night at room temperature with gentle shaking. Non-solubilized material was removed by ultracentrifugation for 90 min at 100,000 *g* and 4°C. The supernatant was subjected to immobilized metal affinity chromatography. To this end, the solubilized material was incubated over night with 2 mL of Talon $Co^{2+}$-beads (Cytiva) at 4°C with gentle agitation in 25 mM Tris, 100 mM KCl, 1 mM $MgCl_2$, pH 8.0. Resin-bound material was packed into a chromatographic column and washed with 20 mL of the same buffer. Elution of KvAP was achieved by addition of 6 mL elution buffer containing 25 mM Tris, 100 mM KCl, 1 mM $MgCl_2$, 100 mM imidazole, pH 8.0. Eluted fractions were pooled and concentrated using Amicon filters (Merck) with a nominal molecular weight cut-off (MWCO) of 10 kg $mol^{-1}$. To remove unspecifically bound proteins, 200 μL of the concentrate was subjected to SEC on an Äkta Purifier 10 system (Cytiva) equipped with a Superose 6 Increase 10/300 GL column at a steady flow rate of 0.2 mL $min^{-1}$ of the mobile phase (25 mM Tris, 100 mM KCl, 1 mM $MgCl_2$, pH 8.0). The eluate was collected and pooled: 8.5–10 mL (fraction 1), 10–12 mL (fraction 2), 12–14 mL (fraction 3), 14–16 mL (fraction 4), and 16–18 mL (fraction 5). SEC fractions were concentrated using Amicon filters having a nominal MWCO of 10 kg $mol^{-1}$. Samples of each fraction were mixed 3:1 (*v*/*v*) with 4-fold concentrated SDS loading buffer, boiled for 10 min at 95°C, and subjected to SDS-PAGE, Western Blot, and TEM (see following section).

**SDS-PAGE and Western blots**

Boiled samples were loaded onto a NuPAGE 4–12% Bis–Tris gel (Thermo Fisher Scientific) and separated by electrophoresis using a constant voltage of 200 V applied for 40 min at 50 W. After separation, gels were fixed and stained in 10% (*v*/*v*) ethanoic acid, 40% (*v*/*v*) ethanol, and 0.025% (*w*/*v*) Coomassie Brilliant Blue for 15 min and destained with 10% (*v*/*v*) ethanoic acid. For Western blotting, proteins were transferred from an unfixed and unstained gel onto a polyvinylidene difluoride (PVDF) membrane (Carl Roth) for 18 min at 15 V and incubated with 3% (*w*/*v*) albumin fraction V in phosphate-buffered saline (PBS; 137 mM NaCl, 3 mM KCl, 10 mM $Na_2HPO_4$, 2 mM $KH_2PO_4$, pH 7.4) for 1 h at room temperature. The membrane was washed with PBS containing 0.1% (*w*/*v*) Tween-20 (PBS-T) and incubated with a primary His-tag antibody at a dilution of 1:3000 (*v*/*v*) in PBS-T for 1 h. The PVDF membrane was washed with PBS-T and PBS and finally incubated with anti-mouse IgG (Fc-specific) alkaline phosphatase secondary antibody at a dilution of 1:1500 (*v*/*v*) in PBS-T. After washing with PBS, antibodies were visualized by addition of NBT and BCIP. Gels and blots were photographed, and protein bands were analyzed with the public-domain software ImageJ.[7]

**TEM**

EM grids were prepared by loading 5 µL of buffered Glyco-DIBMALPs (50 mM Tris, 200 mM NaCl, pH 7.4) made from (0.01–0.05) g $L^{-1}$ DMPC or POPC onto glow-discharged continuous carbon grids (300 mesh; Quantifoil Micro Tools, Großlöbichau, Germany) or SF162 Cu grids coated with Formvar (Plano, Wetzlar, Germany). Excess liquid was blotted off with a strip of filter paper after 30 s followed by two washing steps and staining with 5 µL of (1–2)% (*w*/*v*) aqueous uranyl acetate solution. Specimens were dried and examined in an EM 900 transmission electron microscope (Carl Zeiss Microscopy, Oberkochen, Germany), and micrographs were recorded with an SM-1k-120 slow-scan charge-coupled device (SSCCD) camera (TRS, Moorenweis, Germany).

**MDS**

For MDS, KvAP was site-specifically labeled at the protein's sole cysteine residue at position 243[6] with the thiol-reactive fluorophore Atto-488 maleimide. For this purpose, KvAP was solubilized with the aid of Glyco-DIBMA, separated from unsolubilized material as described above, and concentrated with Amicon filters having a nominal MWCO of 50 kg $mol^{-1}$. The concentrate was incubated overnight using 3 mL Talon $Co^{2+}$-beads in 25 mM Tris, 100 mM KCl, pH 8, at room temperature under gentle shaking. Resin-bound material was packed into a chromatographic column and washed with 30 mL of the same buffer. Protein-loaded beads were incubated for 45 min in 6 mL of 25 mM Tris, 100 mM KCl, 4 mM TCEP, pH 7.5 and washed with 60 mL of the same buffer. Nanodiscs were incubated in 3 mL of 20 mM HEPES, 100 mM KCl, pH 7.4, supplemented with 60 µM Atto-488 maleimide for 2 h in the dark with gentle shaking to conjugate the dye to KvAP. Unconjugated dye was washed out with 6 mL of 20 mM HEPES, 100 mM KCl, pH 7.4, and KvAP-containing nanodiscs were eluted with 25 mM Tris, 100 mM KCl, 100 mM imidazole, pH 7.5. The eluate was concentrated using Amicon filters having a nominal MWCO of 10 kg $mol^{-1}$. Concentrated samples were purified by SEC as described above, and elution fraction 3 was concentrated again by Amicon filters having a nominal MWCO of 10 kg $mol^{-1}$. The concentrate harboring labeled KvAP in Glyco-DIBMALPs was analyzed by MDS. To this end, 5 µL of the concentrate or, for comparison, 5 µL of 100 µg $L^{-1}$ unconjugated ATTO-488 maleimide in the same buffer was pipetted onto a microfluidic chip and analyzed with the aid of a Fluidity One-W instrument (Fluidic Analytics, Cambridge, UK) kept at 23°C.

### 2. Theoretical background

**Pseudophases in lipid/surfactant mixtures**

We have previously shown[8–11] that the solubilization of LUVs by amphiphilic copolymers can be rationalized in terms of a three-stage model[12] that considers lipid (L) and surfactant (S) molecules in bilayer (b) and micellar (m) phases as well as surfactant monomers in the aqueous (aq) phase. The concentrations of lipid and surfactant, $c_L$ and $c_S$, respectively, determine the presence and abundance of each of these phases. In a lipid/polymer mixture, where the polymer assumes the role of the surfactant, an increase in $c_S$ at given $c_L$ leads to a transition from the vesicular bilayer range to the coexistence range, within which polymer-saturated bilayer vesicles coexist with lipid-saturated nanodiscs. Upon a further increase in $c_S$, the vesicles are completely solubilized and transformed into polymer-bounded nanodiscs. In this interpretation of the three-stage model, nanodiscs take the role of mixed micelles found in conventional lipid/surfactant mixtures.[12,13] The first nanodiscs are formed at a threshold known as the saturation (SAT) boundary, while a second transition designated as the solubilization (SOL) boundary marks the completion of nanodisc formation and the concomitant disappearance of the last vesicular structures.

Plotting the $c_S$ values at the SAT and SOL boundaries against the corresponding $c_L$ values gives rise to two straight lines described by

$$c_S^{SAT}=c_S^{aq,o}+m_P/m_{LS}^{b,SAT}c_L \quad (1)$$

$$c_S^{SOL}=c_S^{aq,o}+m_P/m_{LS}^{m,SOL}c_L \quad (2)$$

The slopes $m_P/m_{LS}^{b,SAT}$ and $m_P/m_{LS}^{m,SOL}$ denote the polymer/lipid mass ratios in vesicular bilayers and nanodiscs at which the vesicles become saturated with polymer and at which solubilization is complete, respectively. Ideally, both lines meet at a common ordinate intercept, $c_S^{aq,o}$, which corresponds to the concentration of "free" polymer in the aqueous phase within the coexistence range. In our previous[8–11] and the present phase diagrams, the ordinate intercepts of the SAT and SOL boundaries are negligibly low, so that the concentration of "active" (i.e., solubilization-competent) polymer in the aqueous phase can be taken as $c_S^{aq,o}$ = 0.

**Phase boundaries from $^{31}$P-NMR**

According to the three-stage model, all phospholipid molecules and, thus, all phosphorus nuclei reside in vesicular membranes as long as the surfactant concentration is lower than or equal to $c_S^{SAT}$ (Eq. 1). In solution-state NMR experiments employing relatively narrow sweep widths, the signal arising from $^{31}$P-nuclei in large, vesicular structures is broadened beyond detection. Thus, the area of the $^{31}$P-NMR peak, $A$, is zero in the absence of solubilized lipid

$$A(c_S < c_S^{SAT}) = 0 \quad (3)$$

Once the polymer concentration exceeds $c_S^{SOL}$ (Eq. 2), all phospholipid molecules are solubilized, and the area under the $^{31}$P-NMR peak amounts to

$$A(c_S^{SOL} < c_S) = f\, c_L \quad (4)$$

where $f$ is the proportionality factor between the concentration of solubilized lipids and the experimentally determined peak area. In general, $f$ depends on the experimental conditions but is constant for a given NMR spectrometer operated using identical instrument settings and acquisition parameters. Within the coexistence range, the peak area is expected to be proportional to the extent of solubilization

$$A(c_S^{SAT} < c_S < c_S^{SOL}) = f\, c_L \frac{c_S - c_S^{SAT}}{c_S^{SOL} - c_S^{SAT}} \quad (5)$$

Here, the last term on the right-hand side reflects the fraction of solubilized lipid as given by the lever rule.[13,14] Pairs of $c_S^{SAT}$ and $c_S^{SOL}$ values at a given lipid concentration were obtained by analyzing the areas derived from the corresponding $^{31}$P-NMR signals in terms of Eqs. 3–5.[8–11] In addition to such local fits considering only one lipid concentration at a time, peak areas measured at four different lipid concentrations were globally fitted with Eqs. 3–5 in order to obtain the best-fit $m_P/m_{LS}^{b,SAT}$ and $m_P/m_{LS}^{m,SOL}$ values. 95% confidence intervals of a global fit, using the areas from all lipid concentrations in one fit, and 95% confidence intervals of local fits, using the areas at each lipid concentration independently, were derived by nonlinear least-squares fitting in Excel spreadsheets, as detailed elsewhere.[1]

**Lipid-transfer kinetics among Glyco-DIBMALPs**

Nanoparticles may exchange lipids through (i) desorption of lipid monomers and interparticle diffusion through the aqueous phase[15–17] and (ii) collisions between two or among more nanoparticles.[18–21] If the size and the shape of the lipid-exchanging particles are identical, the observed rate constant due to diffusional lipid exchange is given by[18,20,21]

$$k_{\mathrm{obs,dif}}\,(c_{\mathrm{L}}) = \frac{k_{\mathrm{dif}} c_{\mathrm{L}}}{c_{\mathrm{L}}^{\circ} + c_{\mathrm{L}}} \tag{6}$$

where $k_{\mathrm{dif}}$ is the diffusional lipid-exchange rate constant and $c_{\mathrm{L}}^{\circ}$ and $c_{\mathrm{L}}$ are the total lipid concentrations initially in the donor and acceptor nanodisc populations, respectively. For collision-dependent lipid transfer between two particles, the observed rate constants amount to

$$k_{\mathrm{obs,col}}(c_{\mathrm{L}}) = k_{\mathrm{col}} c_{\mathrm{L}} \tag{7}$$

where $k_{\mathrm{col}}$ is the second-order rate constant characterizing lipid exchange through binary collisions. If both mechanisms contribute to lipid transfer, the overall observed rate constant is given by the sum of Eqs. (6) and (7)[18,20,21]

$$k_{\mathrm{obs}}\,(c_{\mathrm{L}}) = k_{\mathrm{obs,dif}}\,(c_{\mathrm{L}}) + k_{\mathrm{obs,col}}(c_{\mathrm{L}}) = \frac{k_{\mathrm{dif}} c_{\mathrm{L}}}{c_{\mathrm{L}}^{\circ} + c_{\mathrm{L}}} + k_{\mathrm{col}} c_{\mathrm{L}} \tag{8}$$

**Time-dependent FRET traces**

Mixing labeled and unlabeled Glyco-DIBMALPs leads to a redistribution and dilution of NBD-PE and Rh-PE. This increases the average distance between the fluorophores and, thus, reduces donor quenching. Consequently, the fluorescence emission intensity of NBD-PE at 530 nm increases exponentially with time, *t*, according to

$$F(t) = F_{\infty} + e^{-k_{\mathrm{obs}} t}(F_0 - F_{\infty}) \tag{9}$$

Here, $F(t)$ is the intensity at time $t$ after mixing, and $F_0$ and $F_{\infty}$ are the original and final intensities, respectively. For global data analysis, Eq. (8) was inserted into Eq. (9) to yield

$$F(t) = F_{\infty} + e^{-\left(\frac{k_{\mathrm{dif}} c_{\mathrm{L}}}{c_{\mathrm{L}}^{\circ} + c_{\mathrm{L}}} + k_{\mathrm{col}} c_{\mathrm{L}}\right)t}(F_0 - F_{\infty}) \tag{10}$$

In this equation, $k_{\mathrm{dif}}$ and $k_{\mathrm{col}}$ are global fitting parameters, whereas $F_0$ and $F_{\infty}$ are local (i.e., $c_{\mathrm{L}}$-specific) fitting parameters. Best-fit parameter values and 95% confidence intervals were derived by nonlinear least-squares fitting.[1] Inclusion of third- or higher-order rate constants as found necessary for DIBMALPs[22] did not significantly enhance the fit for lipid exchange among Glyco-DIBMALPs.

### 3. Supporting results

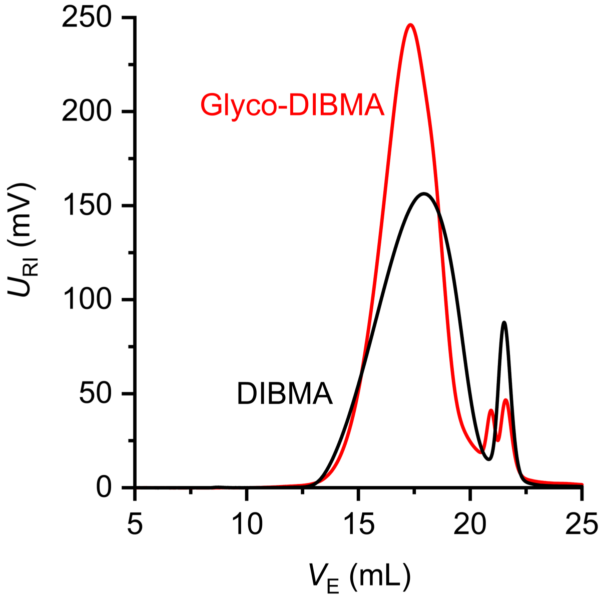


**Supplementary Figure 1:** Analytical SEC of Glyco-DIBMA and DIBMA on a Superose 6 Increase 10/300 GL column in 50 mM Tris, 200 mM NaCl, pH 7.4. Shown is the voltage of the refractive index detector, $U_{RI}$, as a function of elution volume, $V_E$.

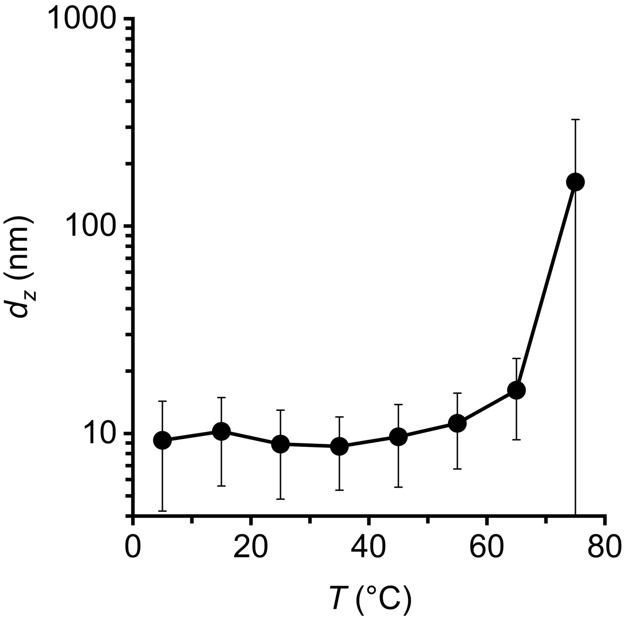


**Supplementary Figure 2:** Colloidal stability of Glyco-DIBMALPs made from POPC as a function of temperature, $T$, in 50 mM Tris, 200 mM NaCl, pH 8.3. Shown are $z$-average hydrodynamic diameters, $d_z$, and corresponding size-distribution widths (“error” bars) of 3.8 g $L^{-1}$ POPC and 3.8 g $L^{-1}$ Glyco-DIBMA.

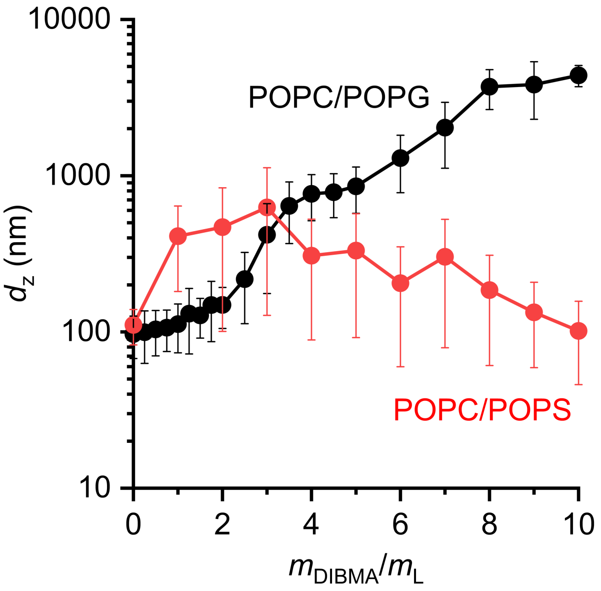


**Supplementary Figure 3:** *z*-Average hydrodynamic diameters and size distribution widths (“error” bars) of LUVs made from POPC/POPG (1:1 mol/mol) or POPC/POPS (7:3 mol/mol), each at ~4 g $L^{-1}$ total lipid, at increasing DIBMA/lipid mass ratios, $m_{DIBMA}/m_L$.
